# Downregulations of mRNA expressions in the *EPAS1* and *EGLN1*: Unraveling a genetic mechanism for high-altitude hypoxia adaptation in Sherpa highlanders

**DOI:** 10.1101/2024.05.23.595497

**Authors:** Yunden Droma, Fengming Yue, Masao Ota, Nobumitsu Kobayashi, Yoshiaki Kitaguchi, Masayuki Hanaoka

**Affiliations:** First Department of Medicine, Shinshu University School of Medicine, Matsumoto, Japan; Department of Histology and Embryology, Shinshu University School of Medicine, Matsumoto, Japan; Department of Medicine, Division of Hepatology and Gastroenterology, Shinshu University School of Medicine, Matsumoto, Japan

## Abstract

Sherpa highlanders exhibit unique hypoxia-tolerant traits, likely evolved through natural selection in response to high-altitude conditions. Previous research has implicated *EPAS1* and *EGLN1* in this adaptation, but their functional mechanisms remain unclear. This study aims to elucidate these mechanisms by exploring mRNA expressions of *EPAS1* and *EGLN1* in Sherpa highlanders. Field investigations enrolled 54 healthy Sherpa highlanders and 25 non-Sherpa lowlanders in Namche Bazaar (3,440 m) and Kathmandu (1,300 m), respectively. Venous blood was collected for erythropoietin (EPO) measurements, *EPAS1* and *EGLN1* mRNA expression analyses, and genotyping of single nucleotide polymorphisms (SNPs) rs13419896 and rs4953354 in *EPAS1*, as well as rs1435166 and rs2153364 SNPs in *EGLN1.* Comparative analyses were conducted on these elements between highlanders and lowlanders. Despite decreased SpO_2_ in Sherpa highlanders, EPO concentrations did not elevate at high altitudes. The *EPAS1* and *EGLN*1 mRNA expressions were significantly lower in Sherpa highlanders compared to non-Sherpa lowlanders. Moreover, genotype distributions and allele frequencies of rs13419896 and rs4953354 in *EPAS1*, as well as rs1435166 and rs2153364 in *EGLN1*, differed significantly between Sherpa highlanders and non-Sherpa lowlanders. Analyses of the relative mRNA expressions of *EPAS1* and *EGLN1* by the SNPs suggested that the rs13419896/G and rs4953354/A alleles in *EPAS1*, as well as the rs1435166/G and rs2153364/A alleles in *EGLN1,* have down-regulatory effects on the gene mRNA expressions in Sherpa highlanders, contributing to inhibiting the EPO hypoxia response and exhibiting hypoxia-tolerant EPO levels in Sherpa highlanders. Natural selection, driven by high-altitude hypoxia, has encouraged adaptive evolution in Sherpa highlanders, including modifications in *EPAS1* and *EGLN1* genes. This evolutionary process leads to the downregulation of gene mRNA expressions, effectively tempering EPO responses to hypoxia and thereby enabling adaptation to high-altitude environments.

## Introduction

The Sherpas, originating from the Tibetan Plateau, have established permanent settlements in the mountainous regions of the Himalayas. Renowned for their expertise in mountaineering and familiarity with high-altitude terrain, Sherpas serve as esteemed guides for mountaineering and trekking expeditions. They exhibit unique hypoxia-tolerant characteristics at high altitudes, demonstrating a gentle response to hypoxia across various physiological traits [1]. Despite the hypoxic conditions, Sherpas at high altitudes do not show significant elevations in hemoglobin (Hb) and erythropoietin (EPO) concentrations, thus preventing chronic mountain sickness (CMS) [2,3]. These hypoxia-tolerant traits are believed to have evolved through natural selection over many generations, driven by environmental hypoxic pressure, enabling adaptation to high-altitude hypoxia [4,5].

Population genomic surveys consistently highlight the genes of endothelial PAS domain protein 1 (*EPAS1*) and egl-9 family hypoxia-inducible factor 1 (*EGLN1*) as potential contributors to high-altitude adaptation in native Tibetans and Sherpas [2,6–8]. Previous studies have demonstrated significant differences in the genotype distributions and allele frequencies of the *EPAS1* and *EGLN1* between Sherpa highlanders and non-Sherpa lowlander controls [2,9]. These significant alterations likely play a role in shaping their hypoxia-tolerant traits by regulating the hypoxia-inducible factor (HIF) pathway in Sherpas at high altitudes [1,3].

These two genes play crucial roles in the HIF signaling pathway, which orchestrates the transcriptional response to hypoxia in humans and other vertebrates [4–6,8,10]. The HIF protein is a heterodimer composed of an oxygen-sensitive α-subunit and an oxygen-insensitive β-subunit. The α-subunit contains an oxygen-dependent degradation (ODD) domain that undergoes hydroxylation by proline-hydroxylase-2 (PHD-2) in the presence of oxygen, leading to proteasomal degradation of the α-subunit under normoxic conditions [11]. However, in a hypoxic environment, the unhydroxylated ODD of the α-subunit prevents proteasomal degradation, resulting in the accumulation of HIF-α subunits. The increased α-subunits heterodimerize with the β-subunits, leading to upregulated HIF and subsequent transcriptional activations of HIF downstream genes, including the gene encoding EPO. Within the HIF pathway, *EPAS1* encodes the HIF-2α subunit, while *EGLN1* encodes the PHD-2 factor [12].

However, the functional mechanisms of the genetic variations in human adaptation to high-altitude hypoxia remain undefined. The identified genetic variations in the *EPAS1* and *EGLN1* associated with high-altitude adaptation are predominantly intronic alleles or located outside the gene boundaries. This suggests that the phenotypic effects could be realized through the regulation of gene expressions by allelic variants, rather than solely directly resulting from mutations in exons involving the dysfunction of HIF proteins [13]. The present study aims to explore the mRNA expressions of the *EPAS1* and *EGLN1* in Sherpa highlanders to provide a relevant genetic explanation for Sherpa adaptation to high altitude, addressing issues related to genetic variants and mRNA expressions of the *EPAS1* and *EGLN1* in the HIF pathway specifically in Sherpa highlanders.

## Materials and methods

The present human genetic study was approved by the Ethics Committee of Review Board of Shinshu University (Matsumoto, Japan, Appended Form No. 2) and the Executive Ethical Review Board of Nepal Health Research Council (Kathmandu, Nepal, 2061-5-28). The protocol of the investigation adhered to the principles outlined in the Declaration of Helsinki of the World Medical Association [14] and written informed consent in the Nepali language was obtained from all participants prior to the study.

### Participants and settings

We enrolled 54 unrelated Sherpa highlanders at an altitude of 3,440 meters (m) for the current study from September 16, 2004, to September 18, 2004, in Namche Bazaar, Nepal [2]. These Sherpa highlanders had been residing in Namche Bazaar since birth, with none reporting symptoms of CMS. Among them, 25 (44.6%) Sherpas worked as trekking guides or porters, frequently ascending to altitudes higher than their residence. Notably, six Sherpa men had successfully reached the summit of Mt. Everest (8,848 m) multiple times. All participants had not traveled to higher or lower altitudes within three months before the investigation. Peripheral oxygen saturation (SpO_2_) and pulse rate were measured using a pulse oximeter (Pulsox-3, Minolta, Osaka, Japan) in Sherpas at 3,440 m. Simultaneously, venous blood samples were obtained from the Sherpas at 3,440 m and promptly transferred into two vacuum blood collection tubes. One of the tubes underwent on-site centrifugation at 3,000 rounds per minute for 10 minutes to separate serum, while the other tube, containing ethylenediaminetetraacetic acid (EDTA) anticoagulant reagent, was used for sampling whole blood. All samples were then stored on-site at -20°C.

The field investigation also recruited 25 non-Sherpa lowlanders at an altitude of 1,300 m in the Kathmandu Valley from September 30, 2004, to October 2, 2004, in Nepal, serving as a control population for comparison with Sherpa highlanders [2]. The Kathmandu Valley is geographically close to Namche Bazaar village, approximately 140 km away. Most of the non-Sherpa lowlanders in the control group were university students, farmers, and housewives who permanently resided at low altitudes. On-site physical examinations, SpO_2_ measurements, and obtaining blood samples for the non-Sherpa lowlanders were all conducted in the Kathmandu valley following the same protocol used for the Sherpa highlanders in Namche Bazaar village. All samples, including those from the Sherpas and non-Sherpas, were shipped to Japan via air cargo while being maintained in a frozen state after the field investigations.

The procedures for subject recruitment and sample collection in both groups have been thoroughly outlined in our prior publications [2,15–18]. Furthermore, the protocols for field investigation and SNP genotyping in laboratory experiments are accessible as a compilation on protocols.io: https://www.protocols.io/run/recruitments-of-sherpa-highlanders-and-non-sherpa-byhdpt26.

### Measurements of serum EPO concentrations

EPO concentrations were measured in the laboratory at Shinshu University in Matsumoto, Japan situated at an altitude of 600 m. The serum samples collected on-site were utilized for measuring EPO concentrations using the Recombigen EPO RIA Kit (Mitsubishi Chemical Medience Corporation, Tokyo, Japan) via radioimmunoassay, following the manufacturer’s instructions.

### Measurements of mRNA expressions of the *EPAS1* and *EGLN1*

Total RNA was extracted from 250 µl of EDTA-anticoagulated venous blood samples collected on-site using RNAiso Blood solution (Cat# 9112; Takara Bio, Japan) following the manufacturer’s protocol. RNA quantity and quality were assessed using a NanoDrop spectrophotometer (Thermo Fisher Scientific, Waltham, MA, USA). RNA samples with an optical density ratio of 260/280 around 2.0 were considered pure, without significant contamination, and suitable for complementary DNA (cDNA) synthesis.

1 µg of gDNA Eraser-treated RNA was reverse transcribed into cDNA using the PrimeScrip® RT reagent Kit with gDNA Eraser (Takara Bio, Japan), following the manufacturer’s instructions. Subsequently, mRNA expression was quantified by quantitative real-time PCR (qPCR) using TB Green® Premix Ex Taq™ II (Tli RNaseH Plus) (Takara Bio, Japan) on a Thermal Cycler Dice Real-Time PCR System (Takara Bio, Japan). Details regarding the primers for mRNA expressions of *EPAS1*, *EGLN1*, and the housekeeping gene *β-actin*, as well as the PCR conditions, are provided in Table 1. Primers for *EPAS1* mRNA expression were obtained from Li et al. [19], while those for *EGLN1* mRNA expression were sourced from Sharma et al. [20]. The mRNA expressions of *EPAS1* and *EGLN1* were normalized to β-actin mRNA and presented as relative quantitative mRNA expression, calculated using the 2^-ΔΔCT^ formula, where ΔΔCT (cycle threshold) is defined as CT _target gene_ – CT _β-actin gene_.

**Table 1.**
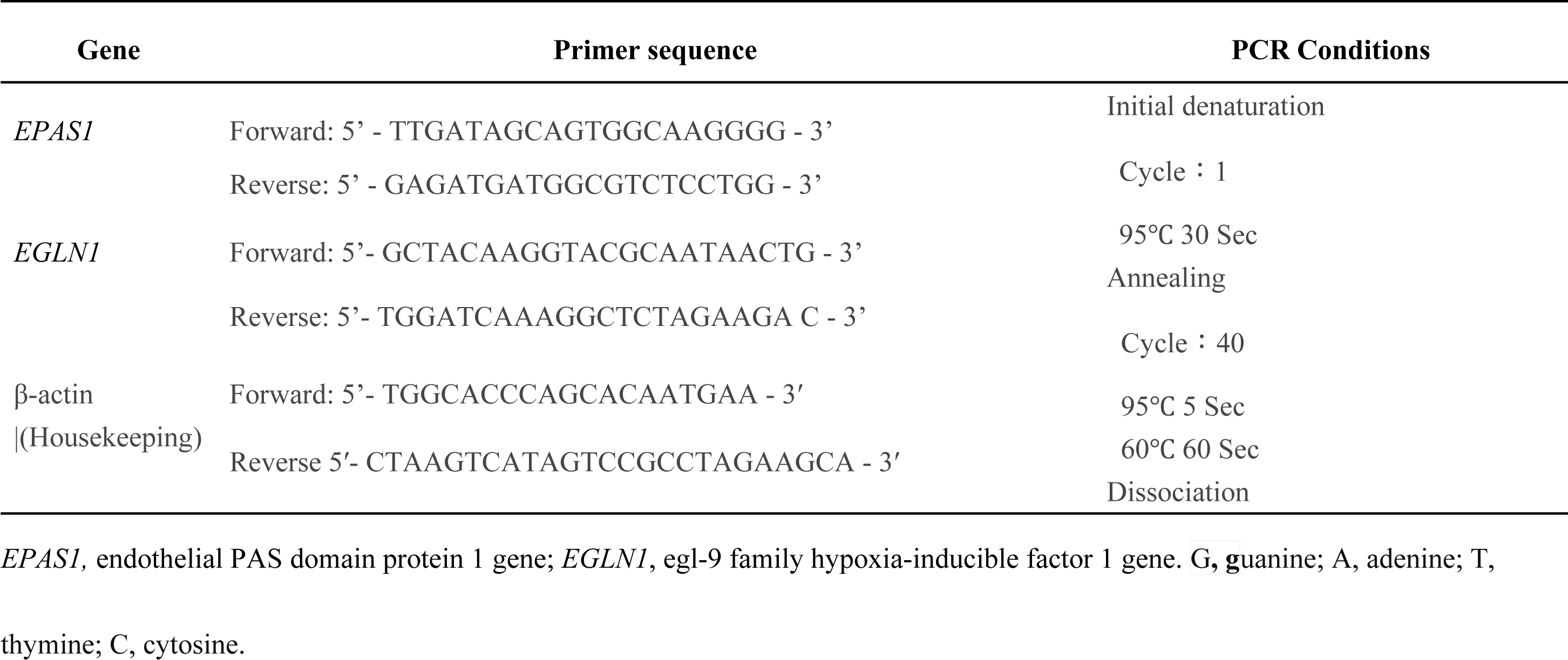
Information regarding the primers for mRNA expression of *EPAS1*, *EGLN1*, and the housekeeping β-actin genes, along with the PCR conditions for quantitative real-time PCR.

### Allele discrimination of the single nucleotide polymorphisms (SNPs) in the *EPAS1* and *EGLN1*

The human *EPAS1* is situated on chromosome 2 and comprises 15 exons spanning 120 kb [11]. The single nucleotide polymorphisms (SNPs) rs13419896 and rs4953354 are located in intron 1 and intron 2, respectively, of the human *EPAS1*. Previous studies have demonstrated a significant association of these SNPs with the lower-than-expected Hb concentrations in Tibetan highlanders residing on the Tibetan Plateau [21]. The human *EGLN1* is positioned on chromosome 1 and consists of 4 coding exons [22]. The SNP rs1435166 is strongly linked pairwise with SNPs in the *EGLN1* that are significantly associated with high-altitude maladaptation in the Indian population [20]. Moreover, the rs2153364 SNP serves as an expression quantitative trait locus (eQTL) for the *EGLN1*, indicating a genomic locus associated with *EGLN1* mRNA expression levels [23]. Information on eQTL is valuable for understanding genetic variations influencing gene expression patterns and can provide insights into the mechanisms underlying various phenotypes [24]. Therefore, the allele discriminations were performed for rs13419896 and rs4953354 in the *EPAS1*, as well as rs1435166 and rs2153364 in the *EGLN1* in the present study. All these SNPs are located within intronic regions.

DNA was extracted from leukocytes in venous blood using the phenol extraction method following established protocols [2]. Allele discriminations for the SNPs in the *EPAS1* and *EGLN1* genes were conducted using the TaqMan® SNP Genotyping Assay with the Applied Biosystems 7500 Fast Real-time PCR System (Applied Biosystems Inc., Foster City, CA, USA), following the manufacturer’s instructions. Genotype data were automatically acquired and analyzed using sequence detection software (SDS v1.3.1; Applied Biosystems, Inc.).

### Statistical analysis

Continuous variables are expressed as mean ± standard error (SE) unless otherwise specified. The differences between the two groups were assessed using Student’s t-test with GraphPad Prism software. The Hardy–Weinberg equilibrium for each SNP was evaluated using exact test procedures (https://wpcalc.com/en/equilibrium-hardy-weinberg/). Differences in genotype distributions and allele frequencies between highlanders and lowlander controls were analyzed using Chi-square tests. P values were corrected (Pc) using Bonferroni’s method for multiple hypotheses testing [25]. The odds ratio and its approximate 95% confidence interval (CI) were calculated. The Pearson correlation coefficient (Pearson’s r) was computed using Microsoft Excel to assess the linear correlation between two variables [26]. Statistical significance was defined as P and Pc values less than 0.05.

## Results

### SpO_2_ and EPO levels in Sherpa highlanders and non-Sherpa lowlanders

There were no significant differences in mean age and gender ratio between Sherpa highlanders (31.3 ± 1.1 years; 23 men/31 women) and non-Sherpa lowlanders (31.7 ± 1.5 years; 13 men/12 women). The SpO_2_ was significantly lower in Sherpas at 3,440 m due to high-altitude hypoxia than in non-Sherpas at 1,300 m (P < 0.0001, Fig 1A). However, despite the decreased SpO2, the serum EPO concentrations did not increase with the hypoxic stimulation in Sherpa highlanders (21.3 ± 1.2 mU/ml), remaining similar to those in non-Sherpa lowlanders (19.4 ± 1.3 mU/ml, P < 0.33, Fig 1B). Additionally, the EPO concentrations were not correlated with the SpO_2_ in Sherpa highlanders (r = -0.24, P > 0.05, Fig 1C), strongly suggesting that EPO exhibits a blunted response to hypoxia in Sherpa highlanders.

**Figure 1:**
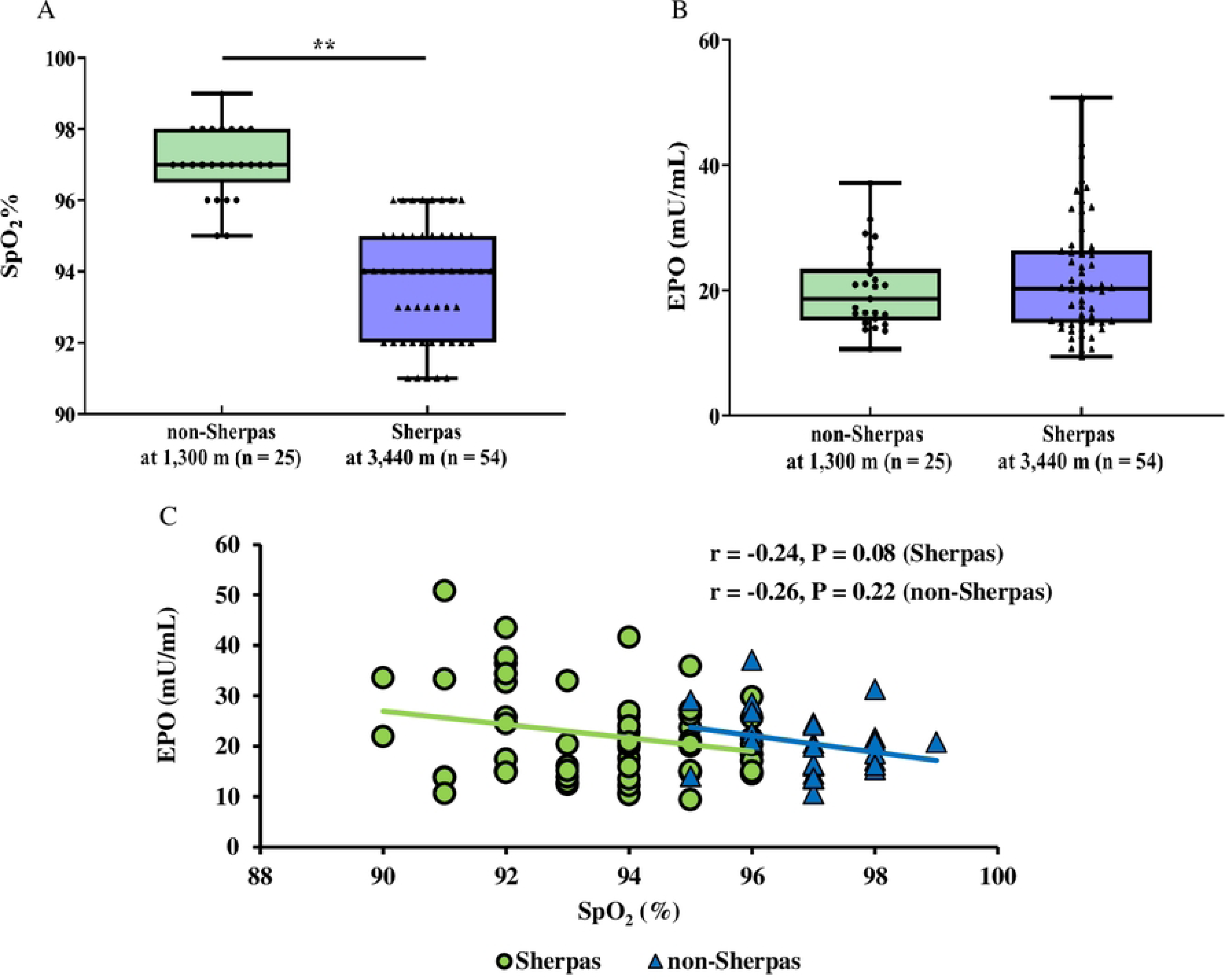
Comparison of phenotypes between Sherpa highlanders at an altitude of 3,440 m (n=54) and non-Sherpa lowlanders at an altitude of 1,300 m (n=25). **A:** Oxygen saturation (SpO_2_%), P = 0.0001; B: Erythropoietin (EPO) concentration, P < 0.33; C: Correlation of SpO2% with EPO concentrations in Sherpa highlanders and non-Sherpa lowlanders. Blue bars represent the Sherpa highlander group; Green bars represent the non-Sherpa lowlander group.

### Relative quantitative mRNA expressions of *EPAS1* and *EGLN1* in Sherpa highlanders versus non-Sherpa lowlanders

The relative quantitative mRNA expressions of the *EPAS1* and *EGLN1* were significantly lower in Sherpa highlanders than in non-Sherpa lowlanders (Table 2, Fig 2A). The *EPAS1* mRNA and *EGLN1* mRNA expressions in Sherpa highlanders were 22.7% and 33.8%, respectively, of the corresponding gene mRNA expressions in non-Sherpa lowlanders, suggesting downregulation of the gene mRNA expressions in Sherpa highlanders relative to non-Sherpa lowlanders. Additionally, the *EPAS1* and *EGLN1* mRNA expressions in Sherpa highlanders were not significantly correlated with the decreased SpO_2_ (Figs 2B and 2C), indicating a blunted response to hypoxia in the gene mRNA expressions in Sherpa highlanders as well.

**Figure 2:**
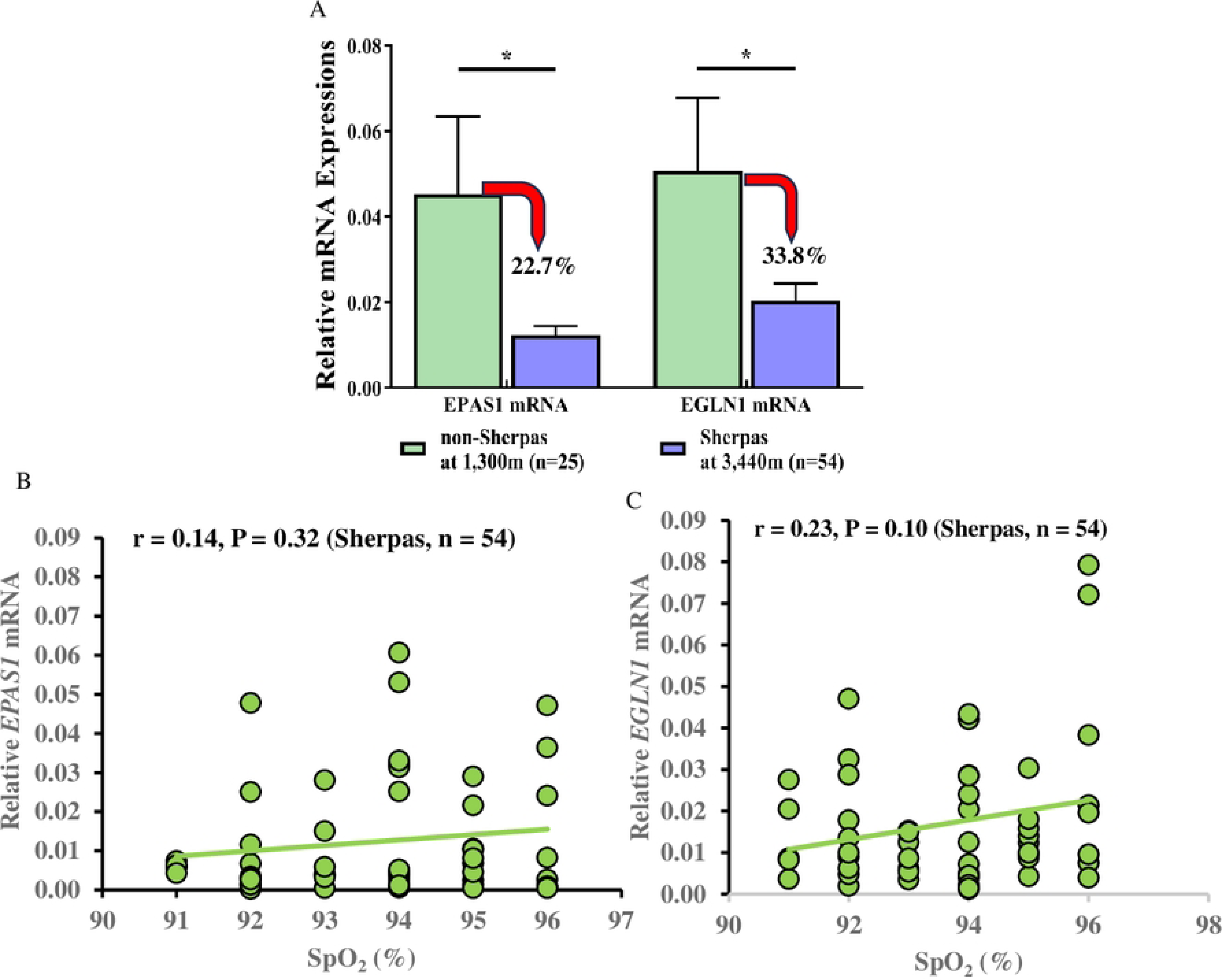
Relative quantitative mRNA expressions of *EPAS1* and *EGLN1* in Sherpa highlanders and non-Sherpa lowlanders. **A:** Comparison of relative quantitative mRNA expressions of *EPAS1* (P = 0.04) and *EGLN1* (P = 0.03) between Sherpa highlanders and non-Sherpa lowlanders; **B:** Correlation of *EPAS1* mRNA expression with SpO_2_% in Sherpa highlanders; **C:** Correlation of *EGLN1* mRNA expression with SpO_2_% in Sherpa highlanders. Blue bars represent the Sherpa highlander group; Green bars represent the non-Sherpa lowlander group; Red arrows indicate the decrease in mRNA expressions in Sherpa highlanders compared to non-Sherpa lowlanders.

**Table 2.** The relative quantitative mRNA expressions of the *EPAS1* and *EGLN1* genes in Sherpa highlanders versus non-Sherpa lowlanders.

| Genes | Relative quantitative mRNA expressions |  | P Value <sup>a</sup> |
| --- | --- | --- | --- |
|  | Non-Sherpa Lowlanders | Sherpa Highlanders |  |
| *EPAS1* | 0.046 ± 0.019 | 0.011 ± 0.002 | 0.04 |
| *EGLN1* | 0.051 ± 0.018 | 0.017 ± 0.002 | 0.03 |

*EPAS1*, endothelial PAS domain protein 1 gene; *EGLN1*, egl-9 family hypoxia-inducible factor 1 gene.
The mRNA expressions of *EPAS1* and *EGLN1* were normalized to β-actin mRNA and presented as relative quantitative mRNA expression, calculated using the $2^{-\Delta\Delta CT}$ formula, where $\Delta\Delta CT$ (cycle threshold) is defined as $CT_{\text{target gene}} - CT_{\beta\text{-actin gene}}$ . The data are expressed as the mean ± standard error.
<sup>a</sup> The differences between the two populations were analyzed using the student's t-test assuming unequal variances.

### Allele discriminations of SNPs in *EPAS1* and *EGLN1* in Sherpa highlanders versus non-Sherpa lowlanders

The genotype distributions and allele frequencies of rs13419896 and rs4953354 in *EPAS1*, as well as rs1435166 and rs2153364 in *EGLN1*, differed significantly between Sherpa highlanders and non-Sherpa lowlanders (Table 3). The frequencies of the major genotypes corresponding to the major alleles of the SNPs rs13419896 and rs4953354 in *EPAS1*, as well as rs1435166 and rs2153364 in *EGLN1*, were significantly reduced in Sherpa highlanders compared to non-Sherpa lowlanders. Conversely, the frequencies of minor genotypes corresponding to the minor alleles of these SNPs were notably increased, resulting in enrichment and becoming the predominant genotypes and alleles in Sherpa highlanders (Table 3). In contemporary Sherpa highlanders, these SNPs displayed entirely different patterns of genotype distributions and allele frequencies compared to non-Sherpa lowlanders, indicating significant genetic divergence among Sherpa highlanders. This divergence is likely the result of natural selection driven by environmental pressures of hypoxia over generations in Sherpa highlanders, facilitating adaptation to high-altitude hypoxia.

**Table 3.**
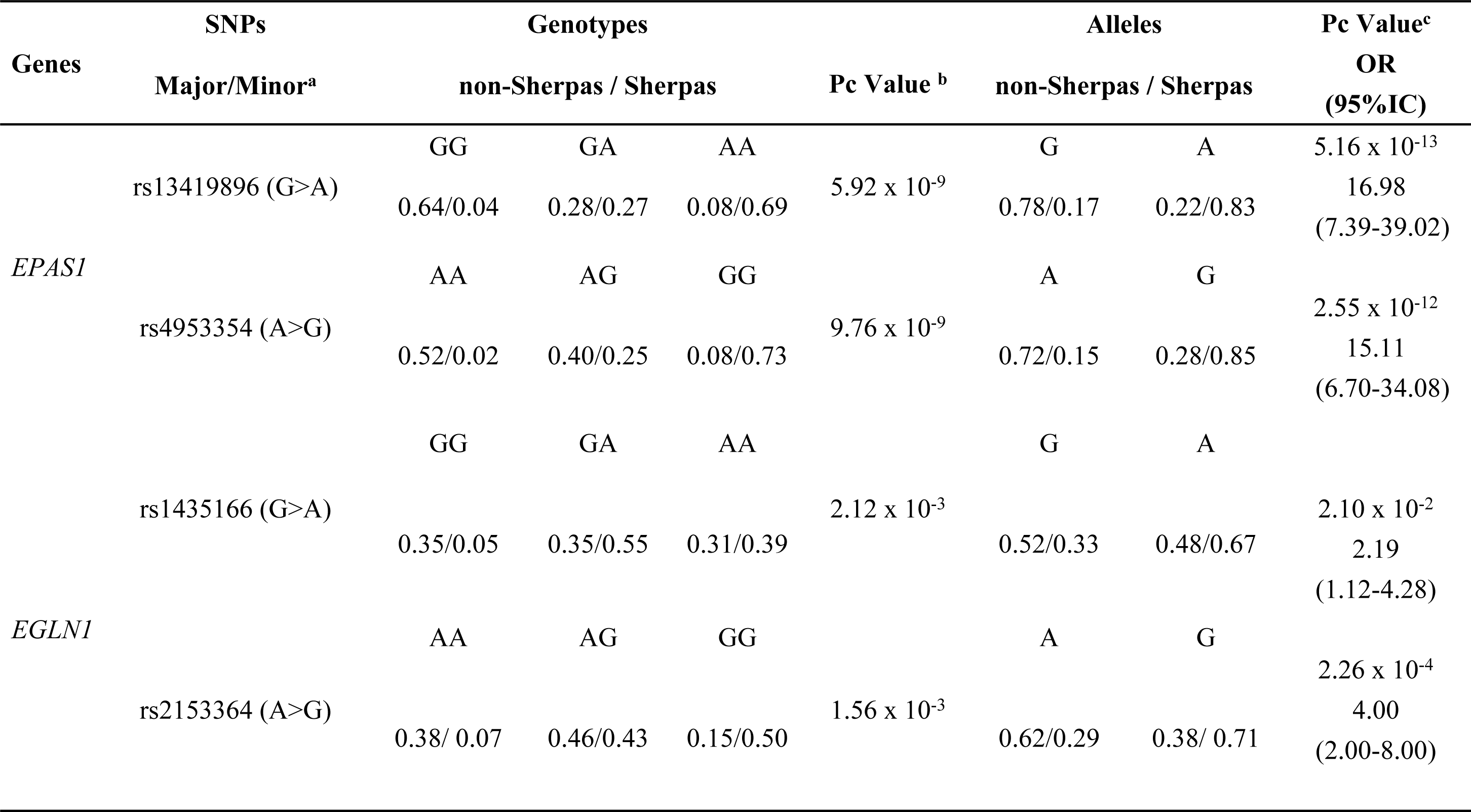
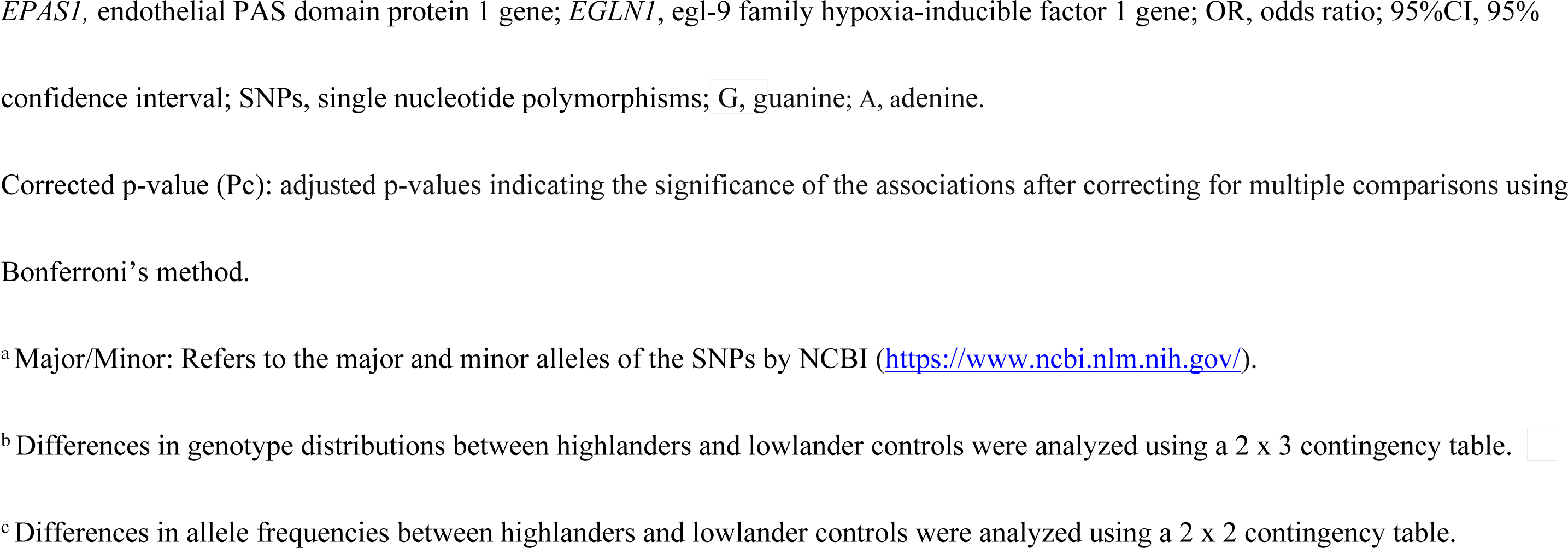
The allele discrimination of the SNPs in *EPAS1* and *EGLN1* in Sherpa highlanders versus non-Sherpa lowlanders.

| Genes | SNPs | Genotypes |  |  | Pc Value<sup>b</sup> | Alleles |  | Pc Value<sup>c</sup> |
| --- | --- | --- | --- | --- | --- | --- | --- | --- |
|  | Major/Minor<sup>a</sup> | non-Sherpas / Sherpas |  |  |  | non-Sherpas / Sherpas |  | OR<br>(95%IC) |
| EPAS1 | rs13419896 (G>A) | GG | GA | AA | 5.92 x 10<sup>-9</sup> | G | A | 5.16 x 10<sup>-13</sup> |
|  |  | 0.64/0.04 | 0.28/0.27 | 0.08/0.69 |  | 0.78/0.17 | 0.22/0.83 | 16.98<br>(7.39-39.02) |
|  | rs4953354 (A>G) | AA | AG | GG | 9.76 x 10<sup>-9</sup> | A | G | 2.55 x 10<sup>-12</sup> |
|  |  | 0.52/0.02 | 0.40/0.25 | 0.08/0.73 |  | 0.72/0.15 | 0.28/0.85 | 15.11<br>(6.70-34.08) |
| EGLN1 | rs1435166 (G>A) | GG | GA | AA | 2.12 x 10<sup>-3</sup> | G | A | 2.10 x 10<sup>-2</sup> |
|  |  | 0.35/0.05 | 0.35/0.55 | 0.31/0.39 |  | 0.52/0.33 | 0.48/0.67 | 2.19<br>(1.12-4.28) |
|  | rs2153364 (A>G) | AA | AG | GG | 1.56 x 10<sup>-3</sup> | A | G | 2.26 x 10<sup>-4</sup> |
|  |  | 0.38/ 0.07 | 0.46/0.43 | 0.15/0.50 |  | 0.62/0.29 | 0.38/ 0.71 | 4.00<br>(2.00-8.00) |

*EPAS1*, endothelial PAS domain protein 1 gene; *EGLN1*, egl-9 family hypoxia-inducible factor 1 gene; OR, odds ratio; 95%CI, 95% confidence interval; SNPs, single nucleotide polymorphisms; G, guanine; A, adenine.
Corrected p-value (P<sub>c</sub>): adjusted p-values indicating the significance of the associations after correcting for multiple comparisons using Bonferroni's method.
<sup>a</sup> Major/Minor: Refers to the major and minor alleles of the SNPs by NCBI (https://www.ncbi.nlm.nih.gov/).
<sup>b</sup> Differences in genotype distributions between highlanders and lowlander controls were analyzed using a 2 x 3 contingency table. ☐
<sup>c</sup> Differences in allele frequencies between highlanders and lowlander controls were analyzed using a 2 x 2 contingency table.

### Relative mRNA expressions of *EPAS1* and *EGLN1* by the SNPs

The *EPAS1* mRNA expressions in Sherpa highlanders identified with the rs13419896/GG (Fig 3A) and rs4953354/AA (Fig 3B) genotypes were 22.1% and 62.9%, respectively, of the *EPAS1* mRNA expressions in non-Sherpa lowlanders with the corresponding genotypes. For *EGLN1*, the mRNA expressions in Sherpa highlanders identified with the rs1435166/GG (Fig 3C), rs1435166/GA (Fig 3C), and rs2153364/AA (Fig 3D) genotypes were 38.1%, 24.5%, and 14.2%, respectively, of the *EGLN1* mRNA expressions in non-Sherpa lowlanders with the corresponding genotypes. Similarly, the *EPAS1* mRNA expressions in Sherpa highlanders carrying the rs13419896/G (Fig 4A) and rs4953354/A (Fig 4B) alleles were 26.4% and 27.7%, respectively, of the *EPAS1* mRNA expressions in non-Sherpa lowlanders carrying the corresponding alleles. The *EGLN1* mRNA expressions in Sherpa highlanders carrying the rs1435166/G (Fig 4C) and rs2153364/A (Fig 4D) alleles were 29.8% and 39.4%, respectively, of the *EGLN1* mRNA expressions in non-Sherpa lowlanders carrying the corresponding alleles. These results suggest that the mRNA expressions of *EPAS1* and *EGLN1* were significantly downregulated in Sherpa highlanders with these genotypes and alleles, likely due to the reduced frequencies of these alleles in Sherpa highlanders (Table 3).

**Figure 3:**
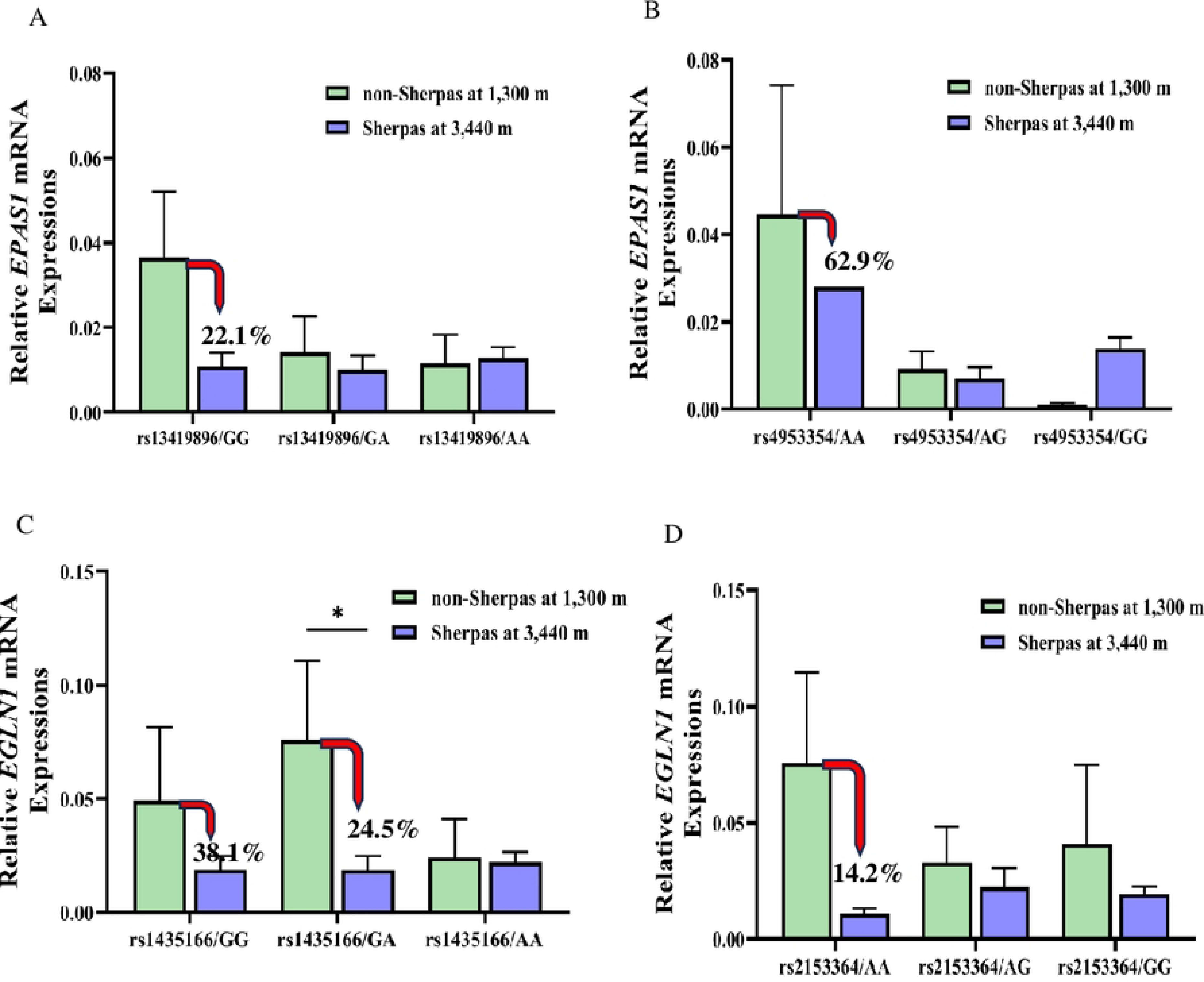
Relative mRNA expressions of *EPAS1* and *EGLN1* by SNP genotypes in Sherpa highlanders and non-Sherpa lowlanders. **A:** *EPAS1* rs13419896 (G>A); **B:** *EPAS1* rs4953354 (A>G); **C:** *EGLN1* rs1435166 (G>A); **D:** E*GLN1* rs2153364 (A>G). Blue bars represent Sherpa highlanders at 3,440 m; Green bars represent non-Sherpa lowlanders at 1,300 m; Red arrows indicate the decrease in mRNA expressions in Sherpa highlanders compared to non-Sherpa lowlanders.

**Figure 4:**
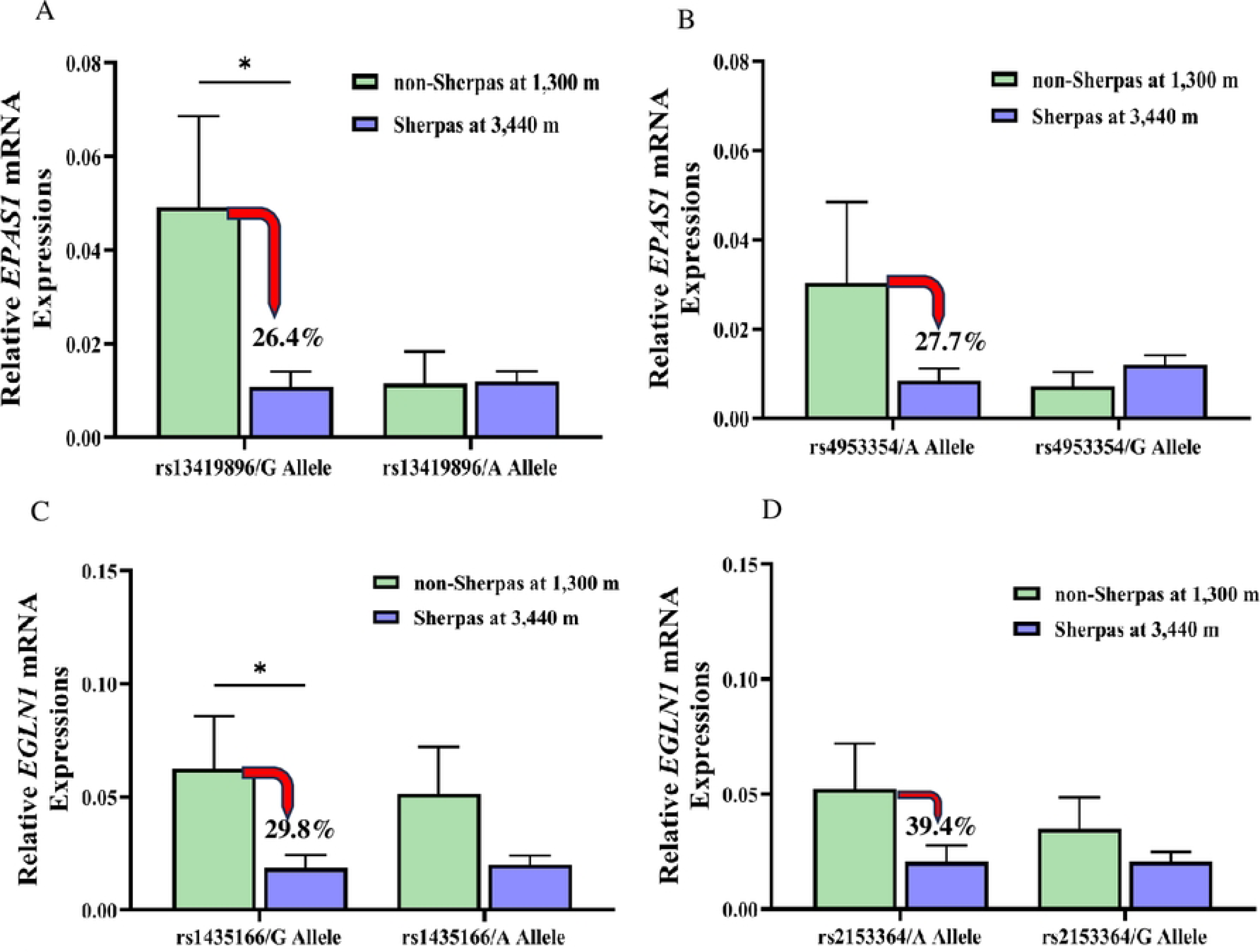
Relative mRNA expressions of *EPAS1* and *EGLN1* by SNP alleles in Sherpa highlanders and non-Sherpa lowlanders. **A:** *EPAS1* rs13419896 (G>A); **B:** *EPAS1* rs4953354 (A>G); **C:** *EGLN1* rs1435166 (G>A); **D:** *EGLN1* rs2153364 (A>G). Blue bars represent Sherpa highlanders at 3,440 m; Green bars represent non-Sherpa lowlanders at 1,300 m; Red arrows indicate the decrease in mRNA expressions in Sherpa highlanders compared to non-Sherpa lowlanders.

### EPO concentrations by the SNPs

There were no significant differences in EPO concentrations between Sherpa highlanders and non-Sherpa lowlanders when categorized by either genotype (Fig 5) or alleles (Fig 6) of these SNPs in *EPAS1* and *EGLN1*.

**Figure 5:**
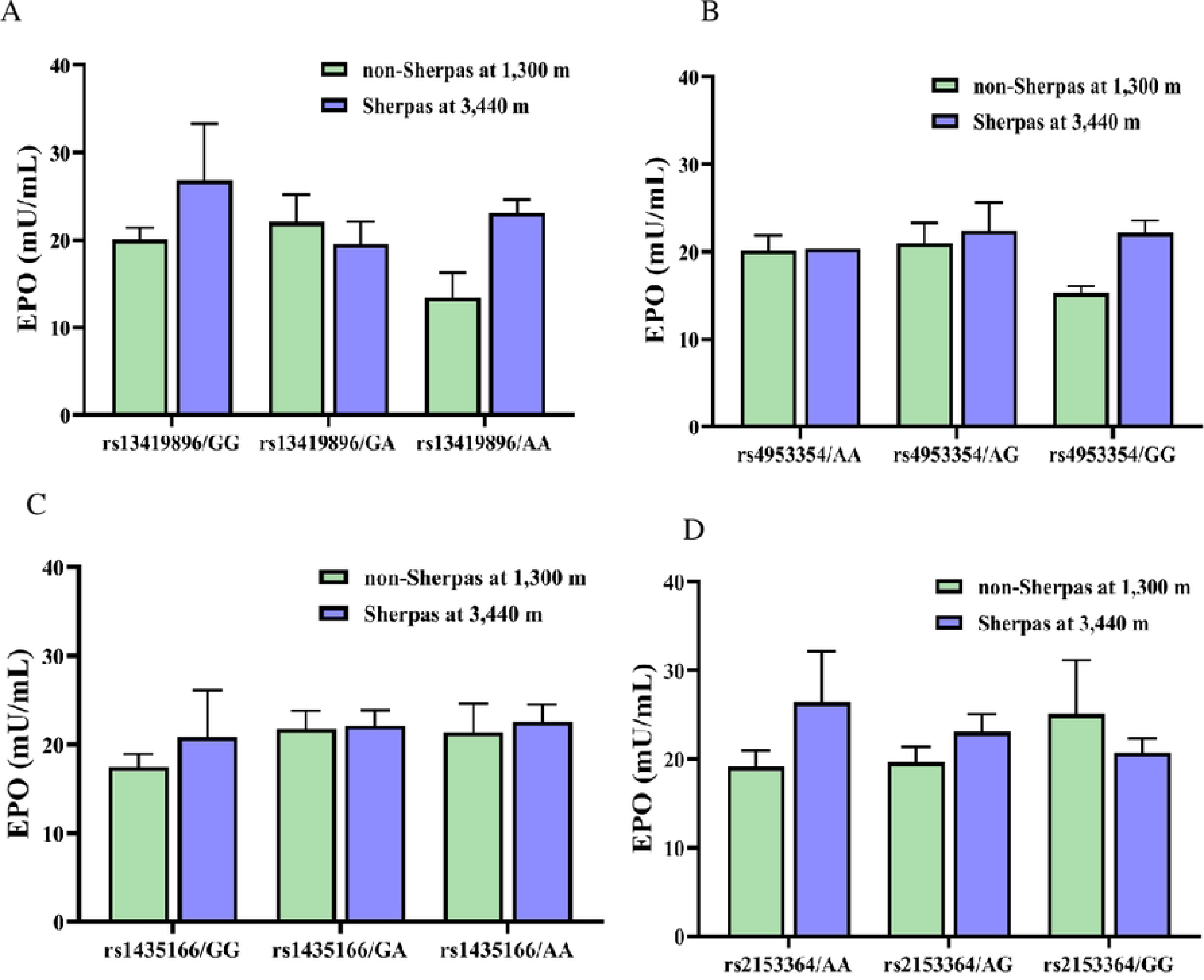
No significant differences in EPO concentrations between Sherpa highlanders and non-Sherpa lowlanders by SNP genotypes. **A:** *EPAS1* rs13419896 (G>A); **B:** *EPAS1* rs4953354 (A>G); **C:** *EGLN1* rs1435166 (G>A); **D:** *EGLN1* rs2153364 (A>G). Blue bars represent Sherpa highlanders at 3,440 m; Green bars represent non-Sherpa lowlanders at 1,300 m; Red arrows indicate the decrease in mRNA expressions in Sherpa highlanders compared to non-Sherpa lowlanders.

**Figure 6:**
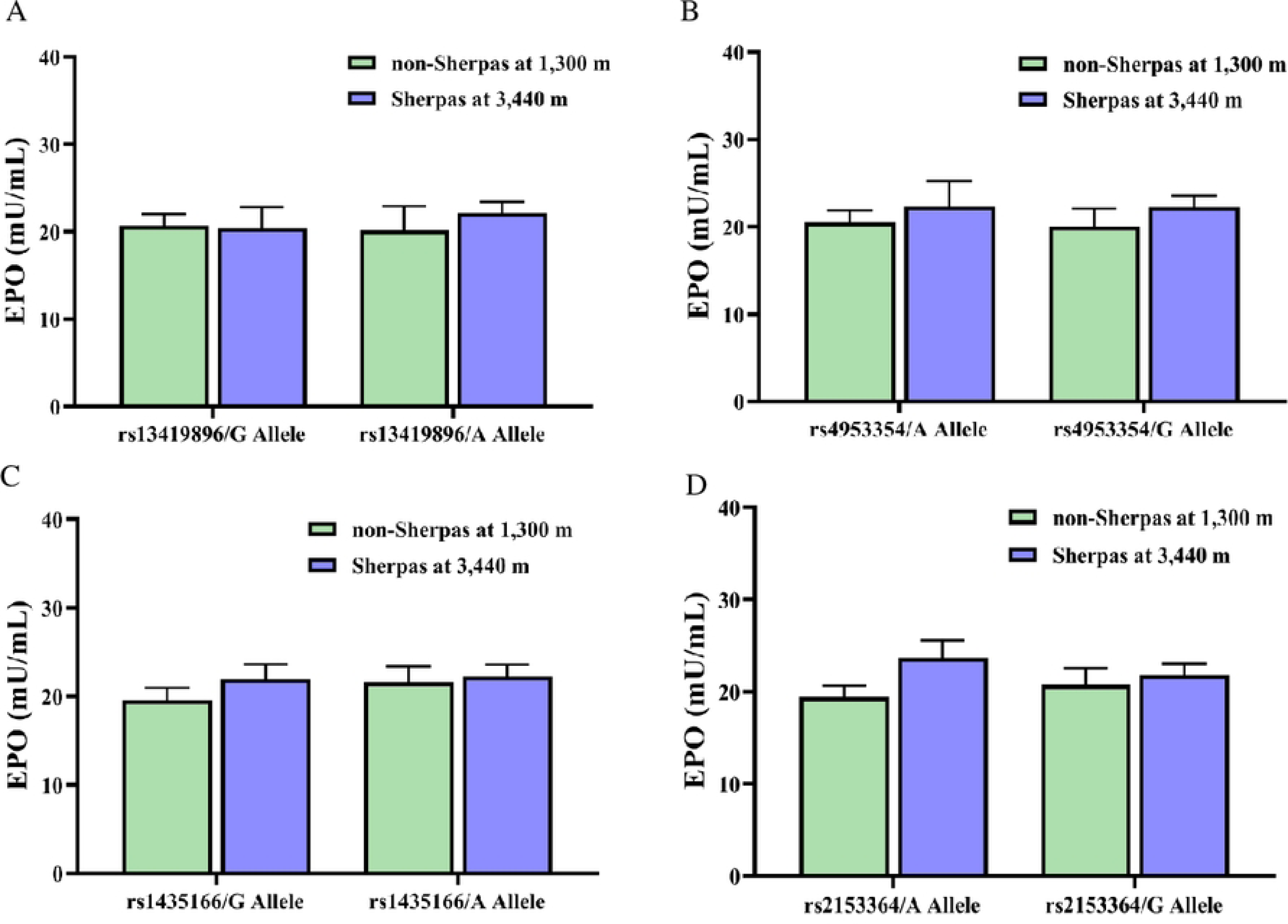
No significant differences in EPO concentrations between Sherpa highlanders and non-Sherpa lowlanders by SNP alleles. **A:** *EPAS1* rs13419896 (G>A); **B:** *EPAS1* rs4953354 (A>G); **C:** *EGLN1* rs1435166 (G>A); **D:** *EGLN1* rs2153364 (A>G). Blue bars represent Sherpa highlanders at 3,440 m; Green bars represent non-Sherpa lowlanders at 1,300 m; Red arrows indicate the decrease in mRNA expressions in Sherpa highlanders compared to non-Sherpa lowlanders.

## Discussion

The initial finding of this study suggests a blunted response of EPO to hypoxia among Sherpas at high altitudes, indicating a hypoxia-tolerant trait in EPO among these native highlanders. A novel discovery reveals downregulation of mRNA expressions of *EPAS1* and *EGLN1* in Sherpa highlanders, likely attributable to the reduced frequencies of rs13419896/G and rs4953354/A alleles in the *EPAS1* gene, and rs1435166/G and rs2153364/A alleles in the *EGLN1* gene compared to those in non-Sherpa lowlanders. Consequently, EPO production in Sherpa highlanders is inhibited at high altitudes, preventing excessive EPO release in response to hypoxic stimulation. As a result, the concentration of EPO in Sherpa highlanders remains within a range typical of individuals living at lower altitudes, facilitating normal physiological processes in Hb and red blood cell (RBC) production. This adaptation aids in coping with high-altitude hypoxia and avoiding CMS.

Erythropoietin (EPO) is a glycoprotein hormone crucial for regulating the production of RBCs and Hb function [27]. At high altitudes, EPO responds physiologically to hypoxia by stimulating the bone marrow to produce more red blood cells, thereby enhancing the oxygen-carrying capacity in circulation and aiding adaptation to high-altitude hypoxia [28]. EPO concentrations significantly increase and peak after 1-2 days, gradually decreasing thereafter but remaining approximately twice the pre-hypoxic values after 10 days of ascending to 4,340 m in healthy lowlanders [29]. However, overproduction of EPO due to hypoxia stimulation can lead to polycythemia, which elevates blood viscosity and increases blood volume, resulting in hematological congestion that eventually accentuates hypoxemia and leads to maladaptation to high altitudes, developing CMS [30]. Blood EPO concentrations are significantly elevated in patients with CMS compared to healthy highlanders [31]. It is noteworthy that EPO concentrations in Sherpa highlanders refuse to increase in response to hypoxia stimulation and exhibit relatively lower-than-expected levels without the effect of decreased SpO_2_ at 3,440 m. This blunted EPO response to hypoxia strongly suggests an inhibition of EPO production in Sherpas at high altitudes, indicating a trait of tolerance to hypoxia involving adaptations to high-altitude hypoxia.

Genome-wide studies consistently identify compelling genetic signatures of natural selection in variants of *EPAS1* and *EGLN1* within the HIF pathway, contributing to high-altitude adaptation in indigenous populations such as Tibetans [7,8,19,21], Sherpas [1–3,9], and Andeans [6]. The present results suggest that the rs13419896/A and rs4953354/G alleles in *EPAS1*, along with rs1435166/A and rs2153364/G alleles in *EGLN1*, were naturally selected, enriching the frequencies of these alleles in Sherpa highlanders for genetic adaptation to environmental hypoxia. This adaptive selection has established a genetic background for the HIF genes that minimize undesirable over-responses to hypoxia in Sherpas living at high altitudes [1,3]. Meanwhile, the frequencies of the rs13419896/G and rs4953354/A major alleles in *EPAS1*, as well as rs1435166/G and rs2153364/A major alleles in *EGLN1* among non-Sherpa lowlanders, degenerated in the Sherpa population during the genetic evolution process. The degeneration of these allele frequencies in the Sherpas led to downregulations of *EPAS1* and *EGLN1* mRNA expressions in Sherpa highlanders, enabling them to live, survive, and reproduce in high-altitude hypoxic environments. Such variant alterations in the intron regions certainly modify gene mRNA expressions in the HIF pathway, providing an advantage for hypoxia-adaptive physiology in Sherpa highlanders.

The transcription factors *EPAS1* and *EGLN1* play pivotal roles as key members of the regulatory network within the HIF pathway [10]. Many *EPAS1* variants are located in noncoding regions, suggesting that they could affect *EPAS1* regulation at the transcriptional level [21,32]. In light of the present results, rs13419896/G and rs4953354/A in *EPAS1*, as well as rs1435166/G and rs2153364/A in *EGLN1,* are major alleles with predominant high frequencies in the lowlander population, determining their quantitative mRNA expressions of *EPAS1* and *EGLN1* (Fig 3 and Fig 4). However, these allele frequencies have significantly diminished, becoming minor alleles in Sherpa highlanders, and play trivial roles in the quantitative mRNA expressions of *EPAS1* and *EGLN1* in the highlanders (Fig 3 and Fig 4). Peng et al. demonstrated that *EPAS1* variants enriched in Tibetans lead to downregulation of *EPAS1* mRNA expression in Tibetan umbilical endothelial cells and placentas. [32]. The downregulated expression of *EPAS1* in Tibetans is associated with relatively low levels of Hb, serving as a protective mechanism against polycythemia and contributing to Tibetan adaptation to high-altitude hypoxia [8,32]. Additionally, Xu et al. observed that native Tibetans possessing wild-type alleles of *EPAS1* variants exhibited attenuated *EPAS1* mRNA expression levels in native Tibetan newborn baby amnions [33]. Moreover, Li and colleagues found that *EPAS1* mRNA expression in native Tibetans at 3,700 meters did not respond to high-altitude hypoxia by increasing gene expression levels [19]. On the other hand, the precise adaptive variants in *EGLN1* involved in high-altitude adaptation remain to be determined [34]. Key controversies persist regarding *EGLN1* gene mRNA expression in native Tibetans and Andeans, including whether it exhibits a gain-of-function phenotype [35] or loss-of-function [36,37] in contributing to high-altitude adaptation [38]. We propose that the significant downregulation of mRNA expressions of *EPAS1* and *EGLN1* is involved in the phenotype of blunted hypoxia response in the hypoxia-sensitive EPO for Sherpas in adaptation to high-altitude hypoxia.

A notable limitation of the current study is the utilization of frozen blood samples for mRNA extraction, which may potentially result in RNA degradation. However, experiments have shown that frozen whole blood stored in EDTA tubes can yield sufficient quality and quantity of total RNA even after long-term storage [39]. The RNAiso Blood reagent (Cat# 9112; Takara Bio) utilized for total RNA extraction in our experiments is designed for rapid and efficient isolation of total RNA from whole blood samples. It operates based on the established acid guanidinium thiocyanate-phenol-chloroform extraction (AGPC) principle for mRNA extraction. In the statistical analysis, to minimize the risk of RNA degradation, gene mRNA expressions were quantified as relative quantitative mRNA expressions, with a β-acting gene mRNA expression serving as the endogenous control, rather than measuring the absolute amount of mRNA expression quantities. Nevertheless, insights into the patterns of gene mRNA expressions between highlanders and lowlanders may offer suggestions for proposing the regulatory mechanisms of the HIF genes in highlander adaptation to high-altitude hypoxia. However, larger sample sizes and the use of fresh blood samples from both highlanders and lowlanders are generally preferred to enhance and verify the present results.

In conclusion, a potential genetic mechanism for high-altitude hypoxia adaptation involves the downregulation of *EPAS1* and *EGLN1* mRNA expressions in Sherpa highlanders. Hypoxia, a natural stressor for humans at high altitudes, has driven the selection of *EPAS1* and *EGLN1* in the HIF pathway, leading to adaptive changes in allele frequencies. These alterations of allele frequencies through the evolutionary process in native highlanders are likely accountable for the downregulations of HIF gene mRNA expressions. Consequently, the downregulated HIF gene mRNA expressions inhibit the sensitive increases of the HIF-associated phenotypes in response to hypoxia, thereby exhibiting tolerance to hypoxia and ultimately enabling adaptation to high-altitude hypoxia.

## Acknowledgments

We express our gratitude to all Sherpa highlanders and non-Sherpa lowlanders for their generous participation in this study. Special thanks to the doctors from the Mountain Medicine Society of Nepal for their invaluable assistance during sample collection in both the Sherpa village and Kathmandu, Nepal. We also appreciate the cooperation extended by the Nepal Health Research Council (Kathmandu, Nepal).

